# Dietary super-doses of cholecalciferol fed to aged laying hens illustrates limitation of 24,25-dihydroxycholecalciferol conversion

**DOI:** 10.1101/2021.10.14.464328

**Authors:** Matthew F. Warren, Pete M. Pitman, Dellila D. Hodgson, Nicholas C. Thompson, Kimberly A. Livingston

## Abstract

**Background:** Older humans who take high levels of vitamin D supplementation for a prolonged time may be at risk of vitamin D toxicity. It is unclear how dietary super-doses (10,000x greater than requirement) can affect vitamin D status in aged animals. Aged laying hens could be a model to compare vitamin D supplementation effects with women in peri-or postmenopausal stages of life.

**Objective:** We investigated dietary super-dose impacts of cholecalciferol (vitamin D_3_) on vitamin D status in aged laying hens in production.

**Methods:** Forty-eight 68 wk old Hy-Line Brown laying hens were individually housed in cages with eight hens per dietary treatment for eleven weeks. Hens were randomly assigned to one of six treatment groups of dietary vitamin D_3_ supplementation and fed *ad libitum*. Supplementation levels were 400, 800, 7,400, 14,000, 20,000, and 36,000 IU D_3_/kg of feed. At the end of the study, all hens were euthanized and tissue samples and feces were collected. Plasma and egg yolk vitamin D metabolites, calcium and phosphorus composition of multiple sites, and tissue gene expression levels were measured.

**Results:** We observed that increasing dietary vitamin D_3_ increased plasma vitamin D_3_ and egg yolk vitamin D_3_ (p < 0.0001 for both sites). We also observed an increase in plasma 24,25-dihydroxycholecalciferol as dietary vitamin D_3_ levels increased (p < 0.0001). The plasma 25-hydroxycholecalciferol:24,25-dihydroxycholecalciferol ratio exhibited an asymptotic relationship starting at the 14,000 IU/kg D_3_ treatment.

**Conclusions:** Dietary super-doses of vitamin D_3_ led to greater plasma and egg yolk vitamin D levels to show that aged laying hens can deposit excess vitamin D in egg yolk. We suggest future research should explore how 24-hydroxylation mechanisms are affected by vitamin D supplementation. Further understanding of 24-hydroxylation can help ascertain ways to reduce risk of vitamin D toxicity.

## Introduction

Older humans taking extremely high levels of vitamin D supplementation for a prolonged time may be at risk of vitamin D toxicity (1, 2). People tend to take vitamin D supplements to increase or maintain vitamin D levels (3). Also, older women take vitamin D supplements to manage the hormonal effects of menopause on bone resorption (4–6). While vitamin D toxicity is uncommon, vitamin D supplements and overfortified foods are the only known means of reaching intoxication levels (1, 7). Vitamin D supplementation can be administered through multiple means with the common routes being oral dose supplementation or dietary supplementation. There was a study that explored how human adults age 65+ given daily oral doses of 1,600 IU cholecalciferol (vitamin D_3_) for 1 year did not exhibit signs of toxicity (8). This illustrates that taking oral doses of vitamin D_3_ in a controlled manner are safe for maintaining vitamin D levels.

An important consideration is if extremely high supplementation of vitamin D_3_ levels over an extended period of time would cause vitamin D toxicity. Chickens have been used extensively to explore questions involving vitamin D_3_ supplementation (9–12). Effects of very high levels of dietary vitamin D_3_ supplementation levels have been explored in chickens over a 48-wk period and the hens did not exhibit histopathological effects related to vitamin D toxicity such as hypercalcemia (10, 11). Considering older humans are likely to take vitamin D_3_ supplements, further exploration of dietary vitamin D_3_ supplementation effects in older hens is necessary. Characterizing how high levels of dietary vitamin D_3_ affect vitamin D metabolism in older hens may help to better understand how overfortified foods can potentially affect vitamin D metabolism in older animals and humans.

Dietary vitamin D supplementation is important for laying hens in production because their bone health is physiologically taxed from egg production (13, 14). Laying hens in commercial farms are fed diets with supplemental vitamin D_3_ beyond the National Research Council (NRC) requirements (15, 16). This ensures the hens can lay eggs and maintain adequate calcium (Ca) absorption for eggshell formation and importantly, bone mineralization (12). There are a few studies which investigated how very high levels dietary vitamin D supplementation affected laying hen production and the metabolic implications pertaining to vitamin D status (10, 11, 17). Altogether, the aforementioned studies illustrate dietary vitamin D_3_ supplementation results in vitamin D-enriched eggs which may be a way to improve vitamin D intake for humans. Further understanding with how very high levels of dietary vitamin D supplementation affects circulating vitamin D metabolite levels in aged laying hens is relevant to the poultry producers interested with extending the production life of laying hens. Also, understanding the impacts of very high dietary vitamin D supplementation in aged laying hens have implications with older women and their vitamin D intake from food fortification.

Our study examined dietary vitamin D_3_ super-dose effects on plasma and egg yolk vitamin D_3_ metabolites and relative gene expression of vitamin D-related genes in aged laying hens in production. We define “super-dose” as treatment doses at least 10,000x greater than requirement. We fed hens diets containing 400, 800, 7,400, 14,000, 20,000, and 36,000 IU D_3_/kg of feed to ascertain vitamin D_3_ supplementation impacts. Hens consuming diets with vitamin D_3_ greater than 7,400 IU D_3_/kg were expected to have increased plasma 24,25-dihydroxycholecalciferol (24,25-(OH)_2_-D_3_) because 24,25-(OH)_2_-D_3_ is an inactive form and would indicate that they reached vitamin D saturation (18–20). Hens consuming super-doses of vitamin D_3_ should also lay eggs with increased vitamin D_3_ content because they would deposit excess vitamin D_3_ into the egg yolk (10).

## Methods

### Animal husbandry

The hens used in our study were from North Carolina State University’s maintained poultry flock. Forty-eight 68-wk old Hy-Line Brown laying hens were housed at North Carolina State University, Raleigh, NC and fed experimental diets for eleven weeks (**Figure 1**). Hens were individually housed in cages between two two-level (top level and bottom level) battery cages with eight hens per treatment. The experimental design was a randomized complete block design with six levels of dietary vitamin D supplementation blocked by cage level. Vitamin D_3_ supplementation levels were formulated to be 250, 500, 1,500, 15,000, 30,000, and 60,000 IU D_3_/kg of feed, but the analyzed vitamin D_3_ levels in the feed were 400, 800, 7,400, 14,000, 20,000, and 36,000 IU D_3_/kg of feed (**Table 1 and Supplementary Table 1**). The source of vitamin D_3_ used in the study was the crystalline vitamin D_3_ from Alfa Aesar (Ward Hill, MA). We refer to the six different analyzed vitamin D_3_ levels as the named treatment groups for our study. In our study, the 400 and 800 IU/kg vitamin D_3_ treatments were formulated to meet the NRC (16) requirements for laying hens. The 14,000, 20,000, and 36,000 IU/kg D_3_ treatments were dietary super-dose treatments for D_3_ intake. Prior to the start of the experiment, all hens were fed the same diet (400 IU D_3_/kg of feed) for one month as a washout period. Hens were fed *ad libitum* along with water. North Carolina State University’s Institutional Animal Care and Use Committee approved all methods for this study, protocol ID number: 18-093-A.

**Figure 1.**
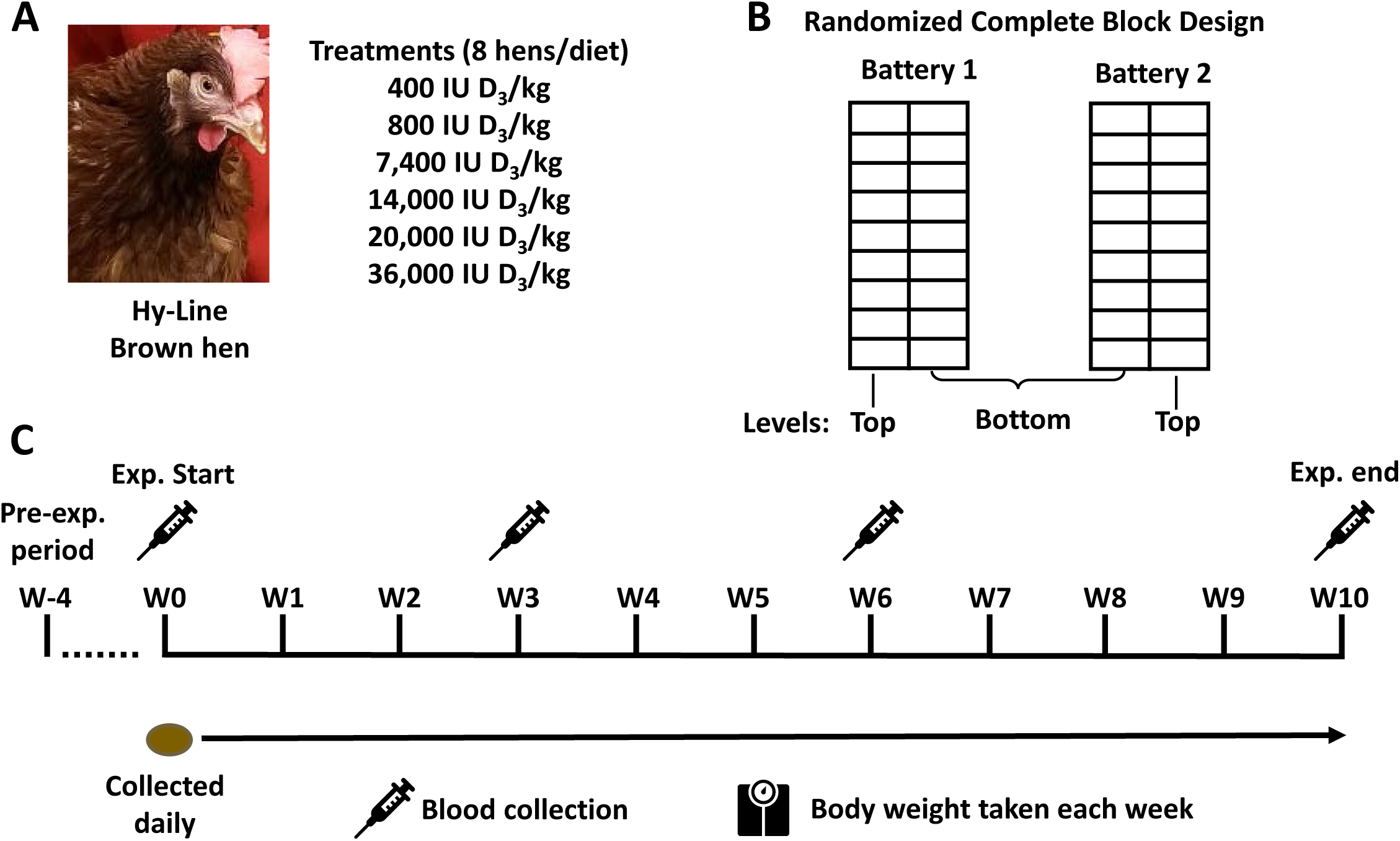
Study design. **A)** Forty-eight 68-wk Hy-Line Brown hens were used in the study and were randomly assorted into one of six treatment groups and fed the diet with the corresponding dietary vitamin D_3_ supplementation level. **B)** Hens were individually housed in cages of a two-leveled battery cage in a randomized complete block design (n = 8/diet). **C)** The experimental timeline in which hens were fed the same basal diet (400 IU D_3_/kg) for four weeks as a washout period (W-4). Hens were started on the experimental diets at week 0 (W0), eggs were collected daily, and weekly body weight was taken until the end of the study. On week 0, 3, 6, and 10, blood was collected from the brachial (wing) vein from all hens to measure ionized blood calcium using an i-STAT™ blood analyzer. Blood was spun down and plasma was collected to measure vitamin D metabolite levels. The study ended on week 10 and all hens were sacrificed and sample tissues were collected.

**Table 1.**
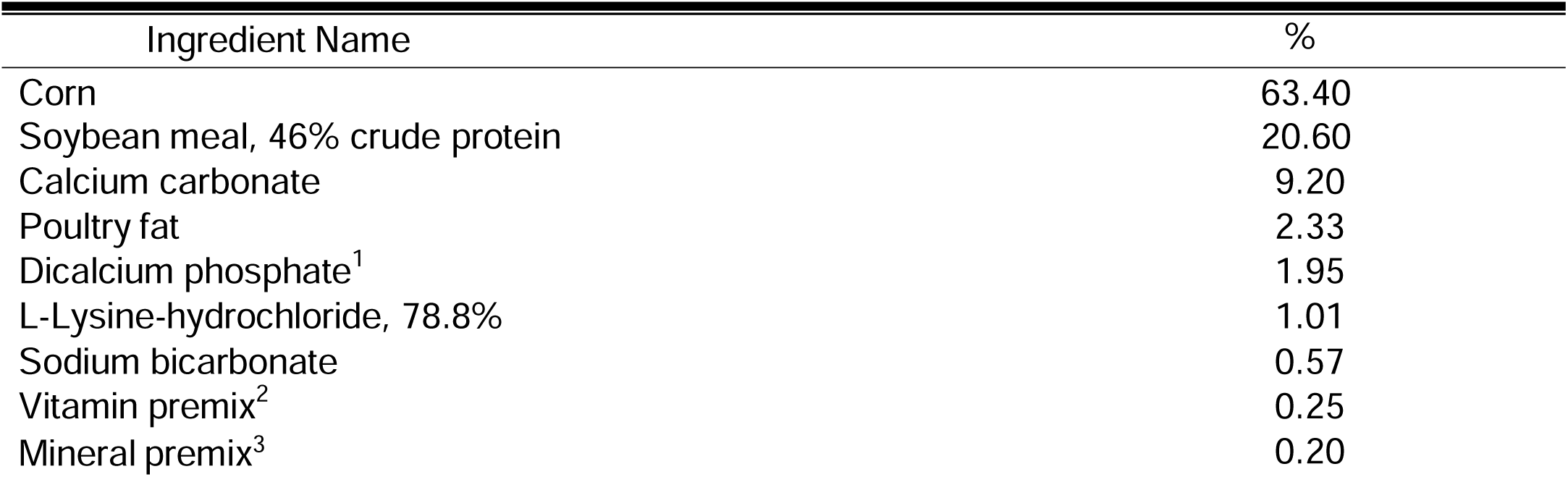
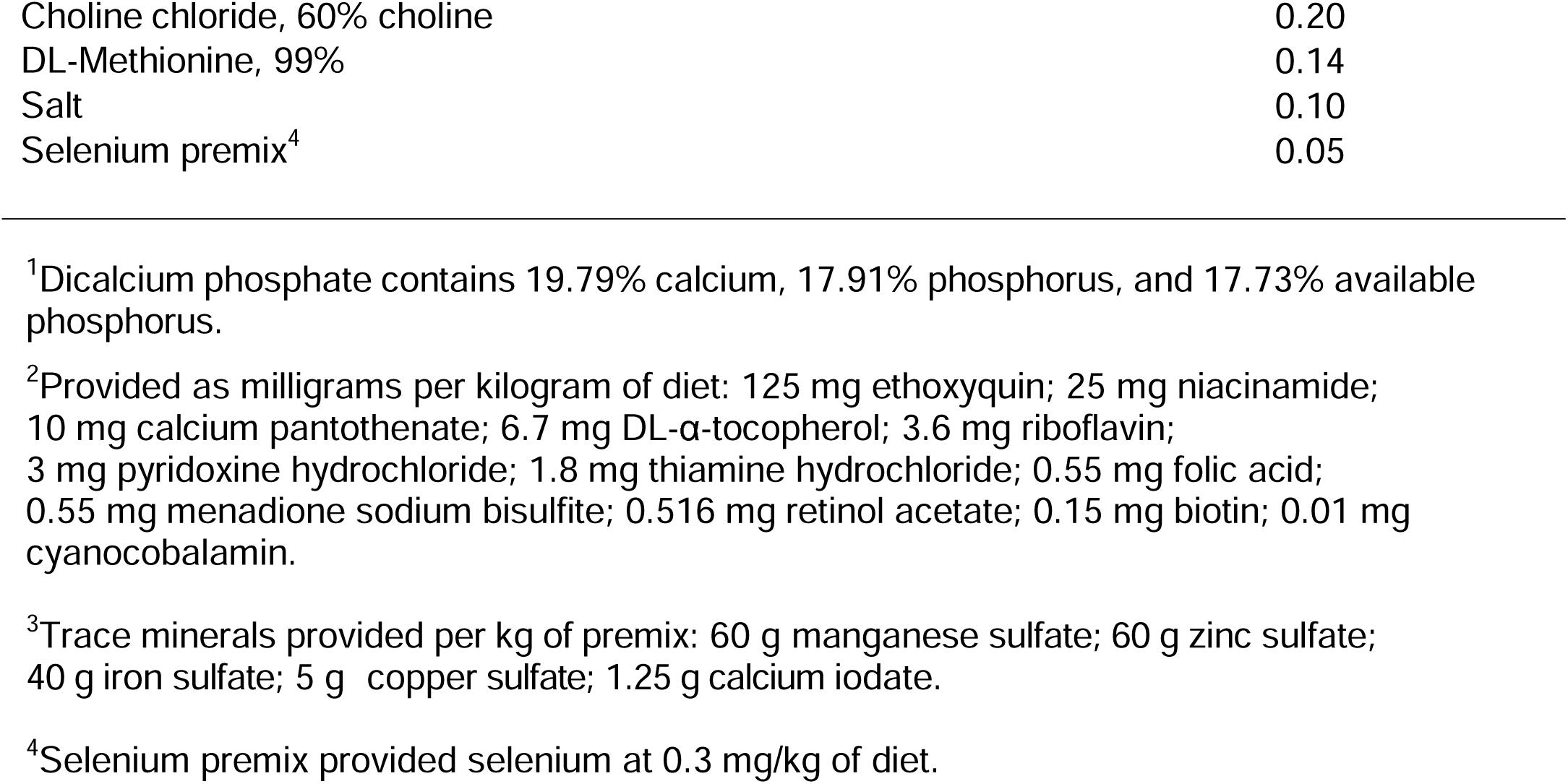
Ingredient composition of the experimental basal diet.

### Sample Collection

Egg collection started 24 hours after the hens were started on the experimental diets. Eggs were collected every morning and stored at 7°C for egg quality analyses. Starting on Mondays, the first egg laid for the week by each hen was selected for egg yolk collection. Eggs were cracked open in a dim-lighted room to reduce photodegradative impacts of light on vitamin D in yolk. The egg yolk was separated from albumen and placed in a small plastic container wrapped in aluminum foil and stored at 4°C for a year until they were freeze-dried using a freeze-dryer (FreeZone 6 Liter Benchtop Freeze Dry System, Labconco, Kansas City, MO). On week 0, 3, 6, and 10, blood was collected from the brachial wing vein from all hens to measure ionized blood Ca using an i-STAT™ blood analyzer (Abaxis, Union City, CA) using CG8+ cartridges (Abaxis, Union City, CA). The remaining blood was spun down to collect plasma which was stored at −80°C. All hens were euthanized by cervical dislocation and tissue samples were collected from 43 hens (minimum of seven hens per treatment) due to time constraints. The duodenum, ileum, liver and kidney were collected and stored in RNAlater at −20°C until RNA extractions were performed and the humerus and tibia bones were collected for measuring Ca and phosphorus (P) composition.

### Calcium and phosphorus content of various sites

Eggshells from weeks 0, 3, 4, 6, and 9 were washed with warm water to remove shell membrane and dried for 48 h at room temperature. Dried eggshells were pre-weighed and further dried at 68°C for 72 h using a dry oven (Blue M, Atlanta, GA) and weighed again. Eggshells were crushed into fine powder and subjected to digestion to measure Ca composition of eggshells. Feces and ileal digesta were also subjected to same steps as eggshells. Dried samples were weighed and then placed in a muffle furnace at 500°C overnight to ash samples. Ashed samples were dehydrated in 2 mL of distilled water and 4 mL of 6 N hydrochloric acid. The resulting sample was mixed and heated to warm to touch. Heated solution was poured into volumetric flask and deionized water was added to 50 mL. The flask was inverted 12 times to mix and the resulting solution was filtered using #40 filter paper into 15 mL centrifuge tubes for analysis. Ca and P were measured by inductively coupled plasma optical emission spectrometry.

The humerus and tibia were wrapped in petroleum ether-moistened cheesecloth and placed in a desiccator for 72 h to extract fat and moisture from the bones. Fat-extracted bones were pre-weighed and dried for 24 h at 100°C to evaporate petroleum ether residues. Fat-and moisture-free bones were weighed and ashed using the same methods as eggshell, feces, and ileal digesta for Ca and P composition and measured by inductively coupled plasma atomic emission spectroscopy.

### RNA extraction and qPCR

Total mRNA was extracted from duodenum, ileum, liver, and kidney using Qiagen’s RNeasy Mini Kit (Germantown, MD). The extracted RNA was diluted and normalized to ∼200 ng/µL for liver and 60 ng/µL for duodenum, ileum, and kidney. The tissues’ RNA was reverse transcribed to complementary DNA (cDNA) using Applied Biosystems’ High-Capacity cDNA Reverse Transcription Kit (ThermoFisher Scientific, Waltham, MA) and their recommended steps to make a 20 µL working solution. Cycling procedure for reverse transcription started with 25°C for 10 min, 37°C for 120 min, 85°C for 5 mins, then held at 5°C indefinitely until storage or use.

Genes amplified for quantitative real time PCR (qPCR) were vitamin D receptor (VDR), 1α-hydroxylase (CYP27C1), 24-hydroxylase (CYP24A1), and glyceraldehyde 3-phosphate dehydrogenase (GAPDH) as the housekeeping gene (**Table 2**). qPCR was conducted using PowerUP SYBR Master Mix (Life Technologies, Grand Island, NY) following manufacturer’s protocol and using Applied Biosystems StepOnePlus Real-Time PCR System (Carlsbad, CA). Cycling procedure started with 95°C for 10 min then 40 cycles of 95°C for 15 s for denaturing and 15 s at 60°C for annealing. All samples were run in triplicates.

**Table 2.**
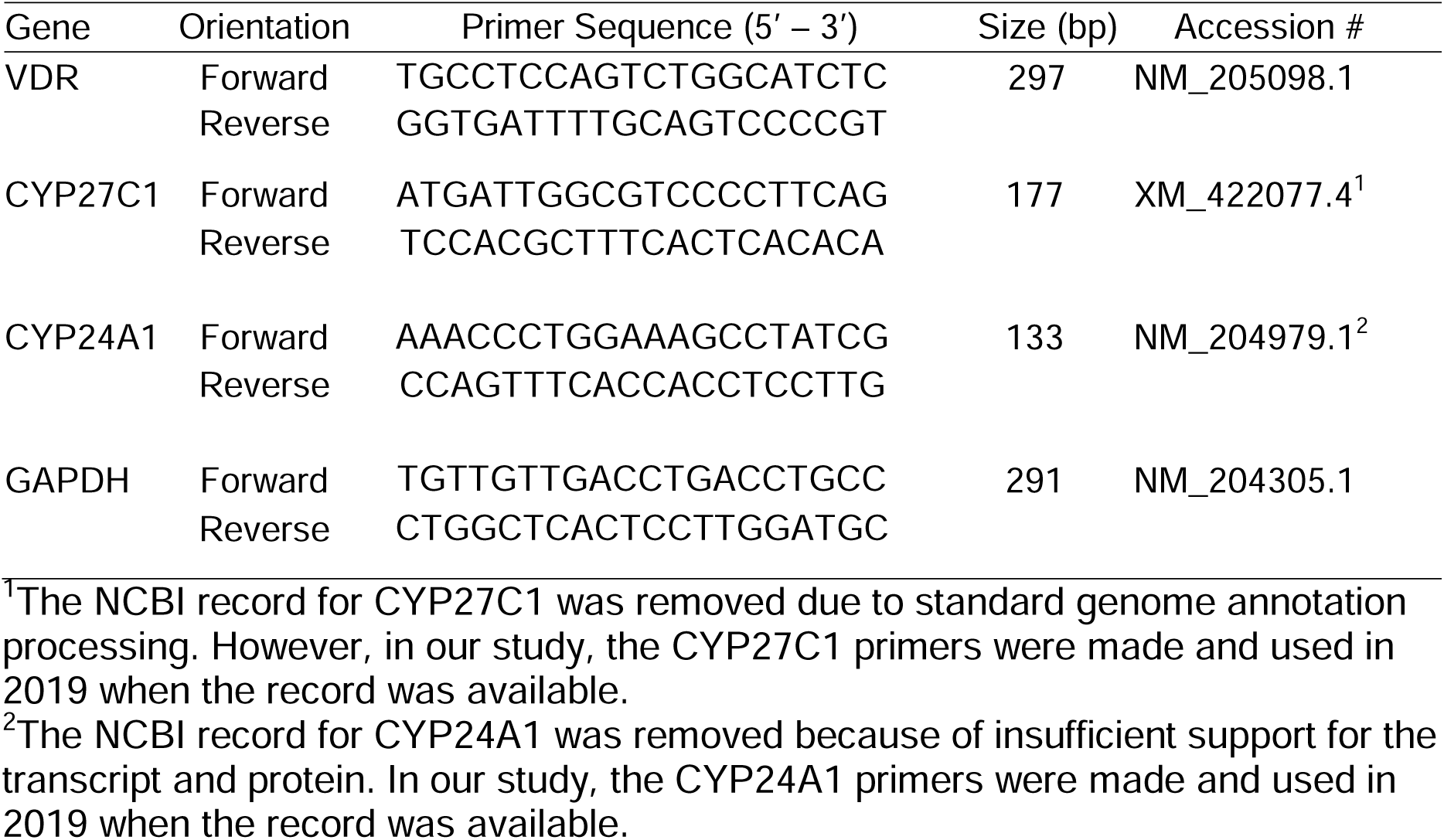
Primer sequences for quantitative real-time PCR.

### Vitamin D metabolites

Plasma from the week 10 timepoint from hens from the 400, 800, 14,000, and 36,000 IU D_3_/kg groups were respectively pooled (n = 4/treatment) and sent to Heartland Assays (Ames, IA) for measuring D_3_, 25-OH-D_3_, and 24,25-dihydroxycholecalciferol [24,25-(OH)_2_-D_3,_ inactive form of vitamin D_3_] using LC-MS/MS. Freeze-dried egg yolk from week 10 timepoint from the same hens were pooled (15 g per sample) like plasma, but only samples from 400, 14,000, and 36,000 IU D_3_/kg groups (n = 4/treatment) were analyzed for D_3_ and 25-OH-D_3_ from Heartland Assays.

### Statistical analysis

We conducted statistical analyses using general linear models using SAS 9.4^®^ for all statistical tests and Tukey-Kramer test was used for multiple comparisons for differences between dietary treatments. We utilized repeated measures to account for temporal effects of weekly body weight, feed intake, egg production, eggshell quality, and ionized blood Ca. Dietary vitamin D_3_ level was the independent variable for Ca and P composition, plasma and egg yolk vitamin D_3_ metabolite concentrations, and gene expression data. Plasma vitamin D_3_ was below the detection limit (< 0.5 ng/mL) for the 400 and 800 IU D_3_/kg treatments so those samples were set to 0.4 as an arbitrary value for statistics to account for model building. All vitamin D_3_ metabolite data exhibited heteroscedasticity and were transformed using natural logarithm function. Transformed data exhibited linear and homoscedastic relationships and were used for statistics. We did not observe a cage level effect in any analysis so the blocking variable was omitted from all statistical tests. All mRNA relative expressions were normalized using 2^-CΔΔT^ with GAPDH as a housekeeper gene. Statistical significance was established at p < 0.05.

## Results

### Hens’ production performance was not influenced by dietary vitamin D_3_

To determine if dietary super-doses of vitamin D_3_ affected the hens’ production value, we measured the hens’ weekly body weights and egg production. Dietary super-doses of vitamin D_3_ did not affect body weight of these laying hens (p = 0.08; **Supplementary Table 2**), but there was an interaction between dietary vitamin D_3_ level over time on feed intake (p < 0.0001). However, there was constant feed wastage throughout the study so this effect could be inflated. A dietary trend was observed for egg production for the entire study duration (p = 0.07; **Supplementary Table 3**). Eggshell strength and eggshell thickness were not affected by dietary vitamin D_3_ levels (p = 0.19, p = 0.72, respectively; **Supplementary Table 3**). There was a trending interaction between dietary vitamin D_3_ level over time in which there was a decrease in eggshell elasticity (p = 0.07).

### Ionized blood calcium is affected by dietary vitamin D_3_ levels

We also examined if ionized blood Ca in the hens was influenced by dietary vitamin D_3_ levels throughout the study. There was no temporal effect on ionized blood Ca (p = 0.65). With week 0 considered as a covariate, there was a dietary effect on ionized blood Ca (p = 0.002, **Supplementary Table 4**). It is not known why there is no clear trend on ionized blood Ca relative to the dietary treatment, but one possibility could be related to when the hen laid an egg prior to the blood collection which may influence circulating Ca levels.

### Fecal calcium is affected by dietary vitamin D_3_ levels, but not ileal digesta or eggshell calcium or phosphorus

We had the eggshell, ileal digesta, and fecal Ca and P measured to determine if dietary vitamin D_3_ fed to our hens would reduce the excretion or loss of Ca and P. There was no dietary effect on eggshell Ca and P (p = 0.64 for both) and ileal digesta Ca or P (p = 0.74 and 0.09, respectively). There was a dietary effect with fecal Ca with hens fed 14,000 IU/kg D_3_ diet had 10.6±0.8% Ca by weight in their feces, whereas all other treatments were 8.0-9.0%, with exception of 36,000 IU/kg D_3_ fed hens which had 7.36±0.43% fecal Ca (p = 0.03, **Supplementary Table 5**). There was no dietary effect on fecal P (p = 0.76).

### Humerus is more calcium and phosphorus dense than tibia

We assessed if dietary vitamin D_3_ would improve bone Ca and P in the hens. However, no dietary effects of vitamin D supplementation were observed on bone Ca or P (p = 0.79 and 0.63, respectively). Humerus bones have more Ca and P than tibia bones (p = 0.020 and 0.015, respectively; **Supplementary** Figure 1).

### Dietary super-dosage levels of vitamin D increased plasma and egg yolk vitamin D_3_ metabolites

We had plasma and egg yolk vitamin D_3_ metabolites measured by LC-MS/MS to determine if dietary vitamin D_3_ affected plasma and egg yolk concentrations. There was a significant increase in plasma concentration of vitamin D_3_, 25-OH-D_3_, and 24,25-(OH)_2_-D_3_ of hens fed dietary super-dose levels of vitamin D_3_ (**D_3_**: p = 0.0002; **25-OH-D_3_**: p < 0.0001; **24,25-OH-D_3_**: p < 0.0001; **Figures 2A-2C**). Although plasma vitamin D_3_ concentration was below limit of detection for the 400 and 800 IU D_3_/kg treatments, the plasma vitamin D_3_ concentration had a strong positive correlation with 25-OH-D_3_ and 24,25-(OH)_2_-D_3_ concentrations (*r =* 0.95, p < 0.0001; *r* = 0.92, p < 0.0001; respectively, data not shown). Plasma vitamin D_3_ and 25-OH-D_3_ had similar concentration values when dietary vitamin D_3_ increased with both metabolites having ∼85 ng/mL at 36,000 IU D_3_/kg treatment. Although plasma 24,25-(OH)_2_-D_3_ concentration was lower than vitamin D_3_ and 25-OH-D_3,_ 24,25-(OH)_2_-D_3_ was affected by dietary treatment and it exhibited a similar rate of increase relative to dietary treatment like vitamin D_3_ and 25-OH-D_3_. The percentage ratio of 24,25-(OH)_2_-D_3_ to 25-OH-D_3_ increased as dietary vitamin D_3_ levels increased and reached an asymptote at the super-dose levels (p < 0.0001; **Figure 2D**). The 24,25-(OH)_2_-D_3_: 25-OH-D_3_ ratio percentage ranged from 8.7% to 20.5% with all super-dose fed hens having a ratio of approximately 20%.

**Figure 2.**
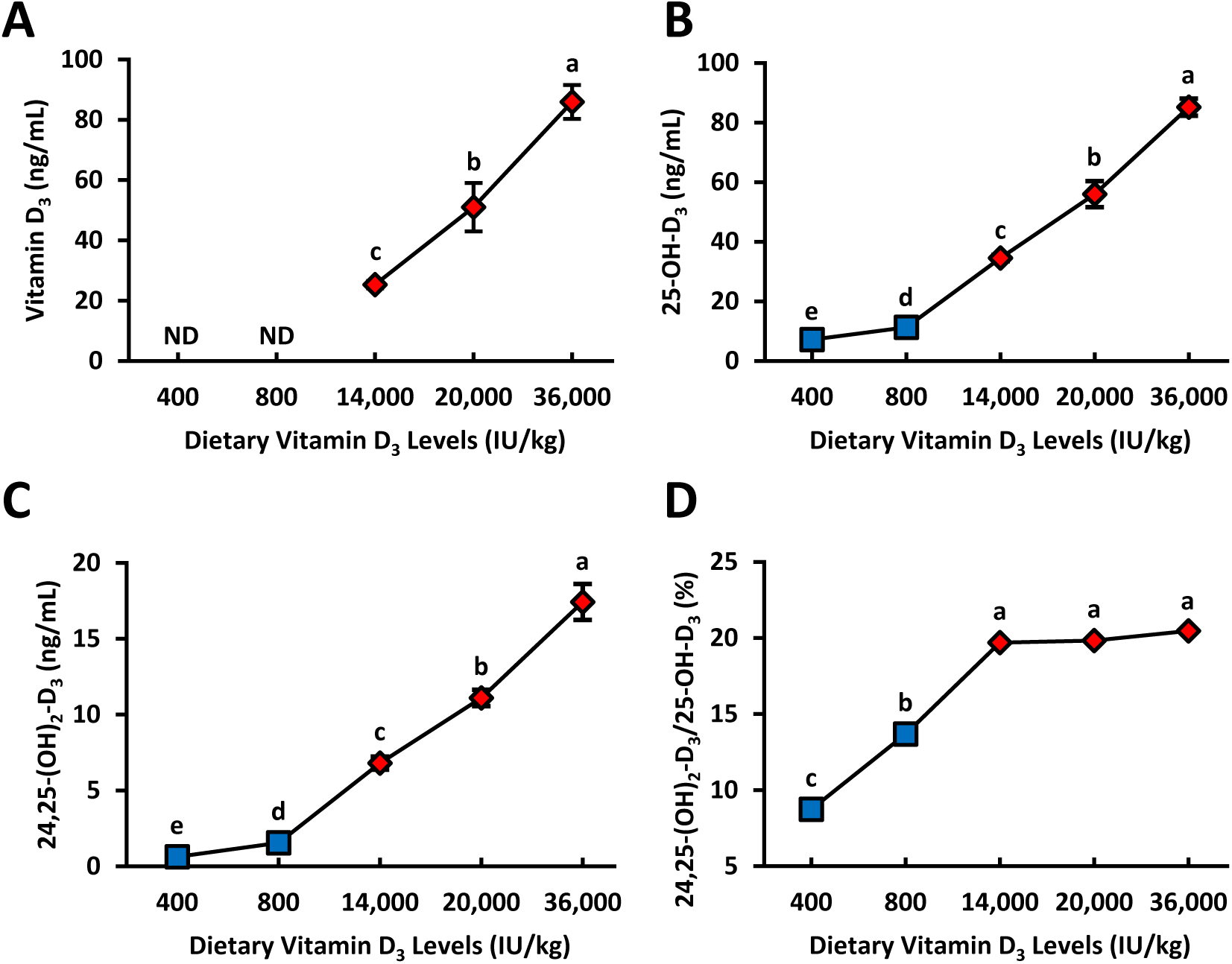
Vitamin D_3_ metabolite plasma concentrations of 78-wk Hy-Line Brown laying hens fed different levels of dietary vitamin D_3_. **A)** Cholecalciferol (Vitamin D_3_; 400 and 800 IU treatment concentrations were not determined) **B)** 25-hydroxycholecalciferol (25-OH-D_3_) **C)** 24,25-dihydroxycholecalciferol (24,25-(OH)_2_-D_3_) **D)** Ratio of 24,25-(OH)_2_-D_3_/25-OH-D_3_ presented as a percentage. Blue squares denote standard NRC range vitamin D_3_ levels (400 and 800 IU D_3_/kg) in diet (n = 4) and red diamonds denote super-dose levels of vitamin D_3_ (14,000, 20,000, and 36,000 IU D_3_/kg) in diet (n = 4). Samples were reported as means ± SEM. Samples with common letters were not statistically different from each other (General linear models, p < 0.0001).

Egg yolk vitamin D_3_ increased drastically as hens’ dietary vitamin D_3_ intake increased (p < 0.0001; **Figures 3A and 3B**). Egg yolk 25-OH-D_3_ was also significantly increased in concentration as hens’ dietary D_3_ increased (p < 0.0001), but the rate of increase was much lower compared to egg yolk vitamin D_3_. Egg yolk vitamin D_3_ was strongly and positively correlated with plasma vitamin D_3_ (*r* = 0.99, p < 0.0001, **Figure 4A**). Egg yolk 25-OH-D_3_ also had a strong positive correlation with plasma 25-OH-D_3_ (*r* = 0.96, p < 0.0001, **Figure 4B**).

**Figure 3.**
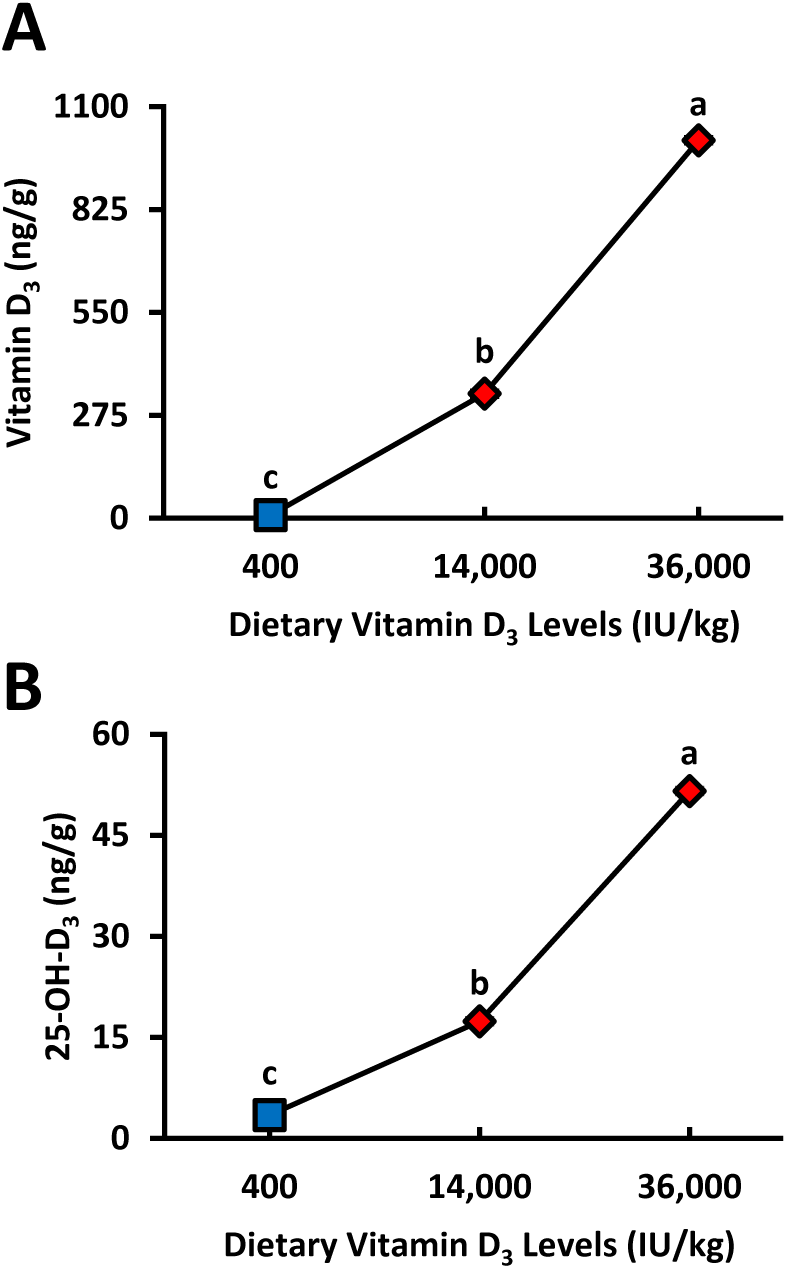
Egg yolk vitamin D_3_ metabolite concentrations from 78-wk Hy-Line Brown laying hens fed different levels of dietary vitamin D_3_. **A)** Cholecalciferol (Vitamin D_3_) **B)** 25-hydroxycholecalciferol (25-OH-D_3_). Blue squares denote standard NRC range vitamin D_3_ levels (400 IU D_3_/kg) in diet (n = 4) and red diamonds denote super-dose levels of vitamin D_3_ (14,000 and 36,000 IU D_3_/kg) in diet (n = 4). Samples were reported as means ± SEM; however, error bars values were narrow and overlapped by the marker for each sample. Samples with common letters were not statistically different from each other (General linear models, p < 0.0001).

**Figure 4.**
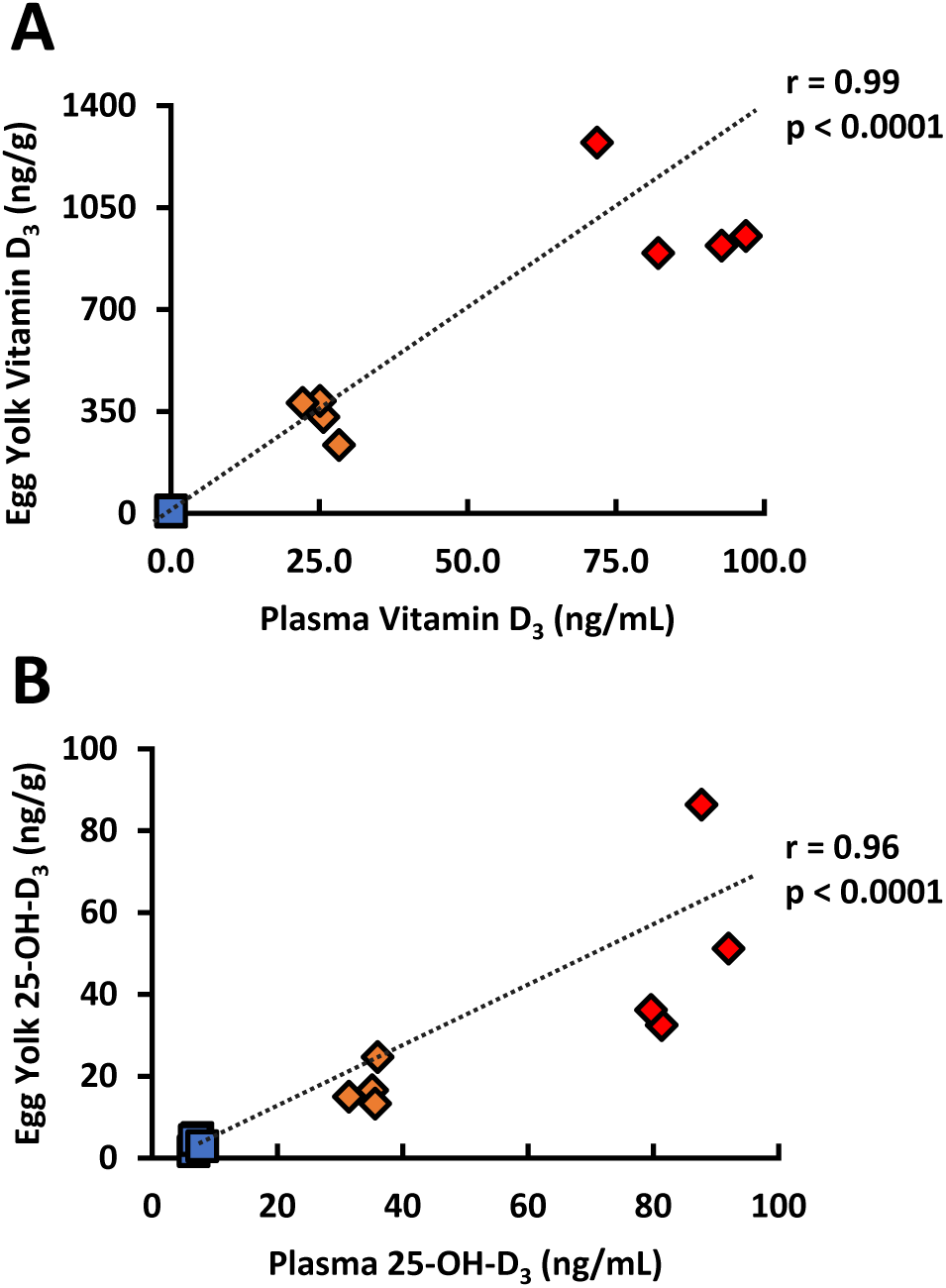
Association between plasma and egg yolk vitamin D_3_ metabolite concentrations from 78-wk Hy-Line Brown laying hens
fed different levels of dietary vitamin D_3_. **A)** Cholecalciferol (Vitamin D_3_) **B)** 25-hydroxycholecalciferol (25-OH-D_3_). Blue squares denote
standard NRC range vitamin D3 level (400 IU D_3_/kg, n = 4) in diet; orange (14,000 IU D_3_/kg, n = 4) and red diamonds (36,000 IU D3/kg, n =
4) denote super-dose levels of vitamin D_3_ in diet (Pearson correlation).

### Dietary super-doses of vitamin D_3_ intake affected vitamin D receptor expression and kidney 24-hydroxylase expression

Considering VDR is the transcription factor responsible for exerting vitamin D’s physiological effects (21), we measured VDR expression in multiple tissues to determine if dietary vitamin D_3_ levels would affect VDR expression. Hens fed higher levels of dietary vitamin D_3_ had upregulated duodenal VDR expression (p = 0.036; **Figure 5A**). There was no dietary effect on VDR expression from ileum, liver, and kidney (p = 0.96, 0.17, 0.32; respectively; **Figures 5B-5D**). We also examined if vitamin D_3_ super-dosages would affect the gene expression of vitamin D hydroxylase enzymes in the kidney. Unexpectedly, kidney 24-OHase expression was lower in hens fed diets with super-dose levels of vitamin D_3_ (p = 0.0006, **Supplementary** Figure 2A). No differences were observed with kidney 1α-OHase expression (p = 0.81, **Supplementary** Figure 2B).

**Figure 5.**
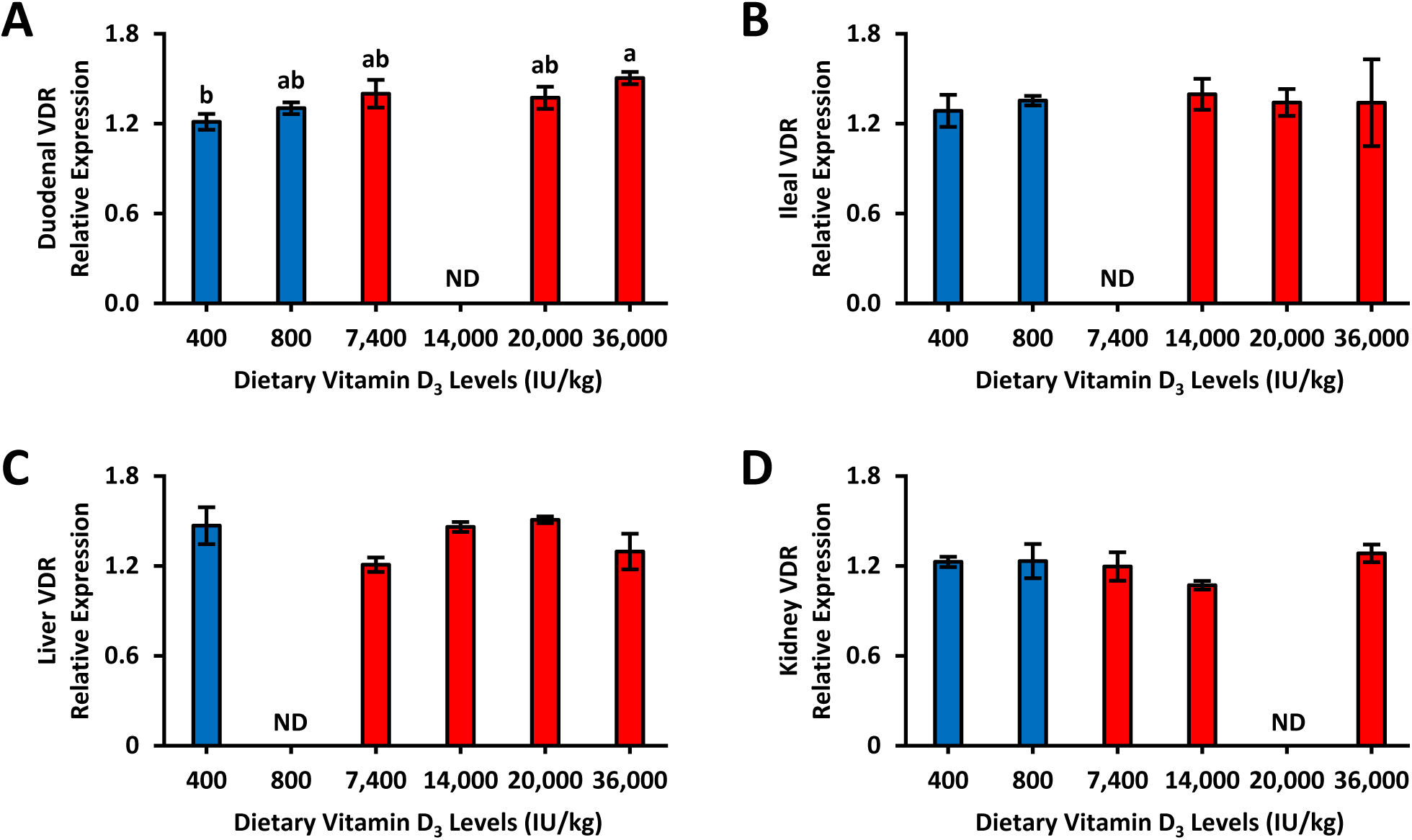
Relative gene expression of vitamin D receptor (VDR) in duodenum, ileum, liver, and kidney of 78-wk Hy-Line Brown laying hens fed different levels of dietary vitamin D_3_. **A)** Duodenal VDR (n = 2-6) **B)** Ileal VDR (n = 2-5) **C)** Liver VDR (n = 2-4) **D)** Kidney VDR (n = 2-4). Tissues were analyzed using qPCR normalized against glyceraldehyde phosphate dehydrogenase (GAPDH; housekeeping gene) expression. Blue bars with dots denote standard NRC range vitamin D_3_ levels in diet and red bars denote super-dose levels of vitamin D_3_ in diet. All samples ran in triplicates and reported as means ± SEM. Bars with common letters were not statistically different from each other (General linear models, p < 0.05).

## Discussion

Our results suggest that dietary super-doses of vitamin D_3_ greatly increased plasma and egg yolk D_3_ levels. Increased plasma vitamin D_3_ indicates these hens absorbed vitamin D_3_ from their diets. Although the inactive vitamin D_3_ metabolite, 24,25-(OH)_2_-D_3_ had a lower measured value than vitamin D_3_ and 25-OH-D_3_, its slope and rate of increase it had the same rate of increase. Increasing plasma 24,25-(OH)_2_-D_3_ concentrations highlights that these hens were likely trying to reduce their circulating vitamin D_3_ levels **(Figure 6)**. In addition, egg yolk vitamin D_3_ drastically increased while yolk 25-OH-D_3_ had a smaller rate of increase. Altogether, there is a strong association between dietary vitamin D_3_ levels and plasma and egg yolk vitamin D_3_ metabolite levels.

**Figure 6.**
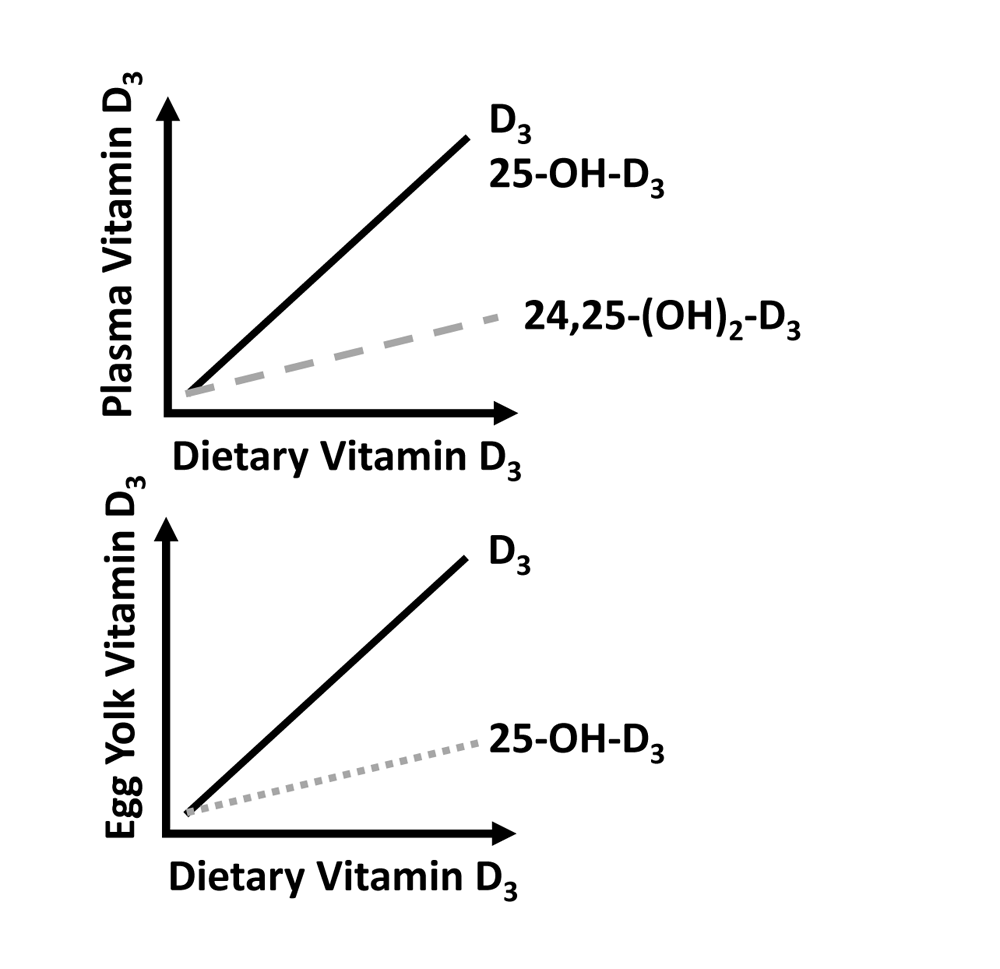
Dietary super-doses of vitamin D_3_ fed to aged laying hens causes drastic increase in plasma and egg yolk vitamin D metabolites. Egg yolk vitamin D_3_ levels were strongly correlated to the dietary levels of vitamin D_3_ fed to the hens. Egg yolk 25-hydroxycholecalciferol (25-OH-D_3_) levels were also dependent on dietary vitamin D_3_; however, 25-OH-D_3_ increased at a lower rate. For plasma vitamin D metabolites, vitamin D_3_ and 25-OH-D_3_ concentrations increased relative to dietary vitamin D_3_ levels fed to the hens. Plasma 24,25-dihydroxycholecalciferol [24,25-(OH)_2_] levels were also dependent on dietary vitamin D_3_, but rate of increase was low.

Although a laying hen’s physiological status affects egg quality (22), high levels of dietary vitamin D_3_ was shown to reduce egg quality, but not affect egg production (23). Mattila et al. (10) and Wen et al. (11) reported that laying hens fed diets with greater levels of dietary vitamin D_3_ throughout their production cycle increased yolk vitamin D_3_ content. The egg production and egg yolk vitamin D_3_ data in our study were similar to the two aforementioned studies, even though the hens in our study were older. A novel finding we observed is that vitamin D_3_ is deposited more readily into yolk compared to 25-OH-D_3_. One possibility is that excess circulating 25-OH-D_3_ was transferred to the egg yolk as a way to lower circulating 25-OH-D_3_ levels. Supplementing laying hen diets with 25-OH-D_3_ increased egg yolk 25-OH-D_3_ and reduced egg yolk vitamin D_3_ (24). It seems likely that the vitamin D metabolite composition in egg yolk is influenced by whatever dietary vitamin D_3_ isoform is fed to the laying hens.

The biological significance of 24,25-(OH)_2_-D_3_ is to reduce 25-OH-D_3_’s plasma concentration (25). To our current knowledge, there is no research on 24,25-(OH)_2_-D_3_ plasma concentration in laying hens. The hens’ plasma 24,25-(OH)_2_-D_3_ increased relative to dietary vitamin D_3_ levels. However, unlike plasma D_3_ and 25-OH-D_3_, the rate of increase with plasma 24,25-(OH)_2_-D_3_ was miniscule. The rate of increase for 24,25-(OH)_2_-D_3_ relative to D_3_ and 25-OH-D_3_ at super-dosage level off at 20%, highlighting a possible asymptotic relationship. The asymptote suggests 24-hydroxylation activity hit its maximal limit. It is not clear if VDR expression is associated with plasma 24,25-(OH)_2_-D_3_ levels or 24-hydroxylation activity because there was little difference in VDR expression across multiple tissues in this study.

Our study has several strengths that highlight its impact with advancing nutritional knowledge. A significant strength of our study is plasma and egg yolk vitamin D metabolite concentration ranges across treatment groups. This illustrates the experimental design captured a broad range of dietary vitamin D_3_ supplementation level effects on plasma vitamin D metabolites that future research can focus on a specific range to build off our findings. Our study provides novel observations of laying hen plasma 25-OH-D_3_ levels relative to dietary vitamin D_3_ supplementation that can be valuable for the poultry industry to consider with vitamin D status. The plasma 24,25-(OH)_2_-D_3_ data is the most exciting finding of our study. Further understanding of the asymptotic relationship of the super-dose levels with plasma 24,25-(OH)_2_-D_3_ concentrations can open new knowledge about 24,25-(OH)_2_-D_3_’s value as a biomarker for vitamin D metabolism.

There were a few limitations with this study that were realized when data was collected. We should have investigated kidney histopathology of these hens because soft-tissue calcification or renal kidney failure could result from the hens reaching vitamin D toxicity (26, 27). However, Mattila et al. (10) did not observe any pathological issues of kidneys from 67-wk old hens fed 15,000 IU D_3_/kg of feed. The smaller sample sizes and missing treatment groups from the qPCR results were because some tissue samples would not yield RNA for cDNA synthesis, even after multiple extraction attempts. It is important to note that the CYP27C1 and CYP24A1 genes may not encode vitamin D-related hydroxylases (28) (Table 1 footnote). We also stored the egg yolk in a refrigerator for about a year before the yolk was freeze-dried; however, our findings are similar to Wen et al. (11) to hint towards minimal vitamin D_3_ degradation (**Table 3**). This could indicate how stable vitamin D_3_ is when it is stored in cold, dark conditions.

**Table 3.** Comparison of egg yolk vitamin D_3_ from eggs laid by hens fed different dietary levels of vitamin D_3_ in this study compared to Wen et al. 2019 study (19)

| **This study<sup>1</sup>** | **Wen et al. 2019 (19)<sup>2</sup>** |
| --- | --- |
| **IU/egg (Dietary treatment IU D<sub>3</sub>/kg)** |  |
| 5.13 (400) | 12.6 (1,681) |
| 199.7 (14,000) | 214.3 (18,348) |
| 605.9 (36,000) | 435.5 (35,014) |

<sup>1</sup>Values were calculated by converting the egg yolk vitamin D<sub>3</sub> concentration (ng/g) to IU and multiplying by 15g (amount used for LC-MS/MS).
<sup>2</sup>These values were reported in Figure 1 for (19).

Our study exhibits that feeding super-doses of dietary vitamin D_3_ to aged laying hens increases their plasma and egg yolk vitamin D_3_. Importantly, there is a possible metabolic limit of 24-hydroxylation to remove excess circulating vitamin D_3_. Investigating 24-hydroxylation mechanisms will be important to understanding vitamin D supplementation impacts in geriatric animals for improving bone health and vitamin D metabolism in older humans.

## Acknowledgments

We thank Gavin Conant, Peter Ferket, Matt Koci, and Shannon Madden for their comments and edits to the initial manuscript; Taylor Jones for her help with the RNA extraction, cDNA synthesis, and qPCR experiments; David Dickey for statistical assistance; Jeff Hall and Zach Spivey for their help with taking care of the hens and collecting samples; Liza Lentz of the Environmental and Agricultural Testing Service laboratory, Department of Crop and Soil Sciences, at North Carolina State University who performed the mineral composition experiments and analyses; and John Rathmacher of Heartland Assays for quantifying the plasma and egg yolk vitamin D_3_. The authors’ responsibilities were as follows-MFW and KAL: designed research; MFW, DDH, NCT, and KAL: conducted research; MFW and KAL: analyzed data; MFW and KAL: wrote manuscript; MFW and KAL: prepared experimental diets for study; MFW and PMP: prepared samples for ashing; MFW and DDH: prepared and shipped plasma and egg yolk samples to Heartland Assays; MFW and KAL: had primary responsibility for final content; and all authors have read and approved the final manuscript.

## Conflict of Interest

All authors report no conflicts of interest.

## Data availability

Data described in the current study will be made available from the corresponding author upon request.

---

**Supplementary Figure 1. Percent by weight of bone calcium and phosphorus from 78-wk Hy-Line Brown laying hens fed different levels of dietary vitamin D_3_. A)** Tibia and humerus calcium **B)** Tibia and humerus phosphorus. Blue squares with lines denote tibia and orange triangles with dashed lines denote humerus (n = 7). Humerus contain more calcium and phosphorus than tibia. Line graphs were expressed as means ± SEM (General linear models, p < 0.05).

**Supplementary Figure 2. Relative gene expression of kidney vitamin D hydroxylases of 78-wk Hy-Line Brown laying hens fed different levels of dietary vitamin D_3_. A)** Kidney 24-hydroxylase (CYP24A1; n = 2-4) **B)** Kidney 1α-hydroxylase (CYP27C1; n = 2-5). Relative expression of vitamin D hydroxylase genes were compared to glyceraldehyde 3-phosphate dehydrogenase (GAPDH; housekeeping gene) expression. Blue bars denote standard NRC vitamin D_3_ level range in diet and red bars denote super-dose levels of vitamin D_3_ in diet. All samples were run in triplicates and reported as means ± SEM. Bars with common letters were not statistically different from each other (General linear models, p < 0.05)

## Notes

### Competing Interest Statement

The authors have declared no competing interest.

### Summary of Updates

Minor change to title.

