## Supplementary Tables for "Dietary super-doses of cholecalciferol fed to aged laying hens illustrates limitation of 24,25-dihydroxycholecalciferol conversion"

**Supplementary Table 1.** Nutrient content of the experimental diet

| Nutrient | % |
| --- | --- |
| Dry matter | 90.46 |
| Moisture | 9.54 |
| Crude protein | 16.00 |
| Fat | 4.98 |
| Fiber | 1.81 |
| Calcium | 4.00 |
| Total phosphorus | 0.67 |
| Ash | 12.44 |
| Available phosphorus | 0.46 |
| Methionine+Cysteine | 0.64 |
| Lysine | 1.57 |
| Methionine | 0.40 |
| Cysteine | 0.26 |
| Lysine | 1.57 |
| Tryptophan | 0.19 |
| Threonine | 0.60 |
| Isoleucine | 0.72 |
| Histidine | 0.43 |
| Valine | 0.83 |
| Leucine | 1.43 |
| Arginine | 1.00 |
| Phenylalanine | 0.83 |
| Metabolizable Energy, kcal/kg | 2,900 |
| Dietary electrolyte balance, mEq/100 g | 202 |

**Supplementary Table 2.** Body weight and feed intake of 78-wk old Hy-Line Brown laying hens fed diets with different levels of vitamin D_3_ over an eleven-week period^1^

| **Treatment (IU Vitamin D_3_/kg of feed)** | **Body Weight (kg)** | **Feed Intake (g/day)** |
| --- | --- | --- |
| **400** | 2.13±0.09 | 128.6±4.9 |
| **800** | 2.32±0.07 | 134.3±3.5 |
| **7,400** | 2.36±0.05 | 167.4±3.6 |
| **14,000** | 2.21±0.10 | 137.3±7.3 |
| **20,000** | 2.18±0.08 | 140.5±8.8 |
| **36,000** | 2.12±0.05 | 171.3±4.4 |
|  | -----------------**P-Value**---------------- | |
| Dietary Vitamin D Level | 0.08 | **<0.0001** |
| Week | **<0.0001** | **<0.0001** |
| Vit D x Week | **0.19** | **<0.0001** |

^1^Means ± SEM were given and p-values were generated by repeated measures general linear model with week as the repeated measurement and the dietary vitamin D_3_ level as the independent variable. Statistical significance established when p < 0.05 (n=8 hens/dietary treatment).

**Supplementary Table 3.** Egg production and eggshell quality measurements of 78-wk old Hy-Line Brown laying hens fed diets with different levels of vitamin D_3_ over an eleven-week period^1^

| **Treatment (IU Vitamin D_3_/kg of feed)** | **Egg Production (%)** | **Shell Strength (kg)** | **Shell Thickness (mm)** | **Shell Elasticity (mm)** |
| --- | --- | --- | --- | --- |
| **400** | 77.4±1.9 | 4.51±0.19 | 0.374±0.010 | 0.209±0.003 |
| **800** | 86.8±1.6 | 4.51±0.10 | 0.371±0.006 | 0.222±0.006 |
| **7,400** | 80.4±3.4 | 4.58±0.17 | 0.372±0.010 | 0.221±0.004 |
| **14,000** | 86.7±2.6 | 4.13±0.11 | 0.360±0.010 | 0.205±0.007 |
| **20,000** | 79.4±3.9 | 4.29±0.12 | 0.366±0.009 | 0.208±0.006 |
| **36,000** | 85.0±1.7 | 4.36±0.14 | 0.377±0.008 | 0.207±0.007 |
|  | ------------------------------------------------**P-Value**----------------------------------------------- | | | |
| Dietary Vitamin D Level | 0.07 | 0.19 | 0.72 | **0.044** |
| Week | ND^2^ | **0.01** | 0.29 | **<0.0001** |
| Vit D x Week | ND | 0.10 | 0.10 | **0.072** |

^1^Egg production was for the entire study period so the p-value was determined by general linear model with dietary vitamin D_3_ as the factor and Tukey’s post-hoc test for multiple comparisons. Eggshell strength, eggshell thickness, and eggshell elasticity were analyzed using repeated measures general linear model with the temporal effect as the repeated measurement and the dietary vitamin D_3_ level as the independent variable. All reported values are means ± SEM. Statistical significance established when p < 0.05 (n=8 hens/dietary treatment).

^2^ND = Not determined

**Supplementary Table 4.** Ionized blood calcium of 78-wk old Hy-Line Brown laying hens fed diets with different levels of vitamin D_3_ over an eleven-week period^1^

| **Treatment (IU Vitamin D_3_/kg of feed)** | **Week 0** | **Week 3** | **Week 6** | **Week 10** | **P-Value** |
| --- | --- | --- | --- | --- | --- |
| **400** | 1.84±0.05 | 1.59±0.08 | 1.68±0.05 | 1.58±0.04 |  |
| **800** | 1.59±0.02 | 1.62±0.07 | 1.69±0.09 | 1.68±0.10 |  |
| **7,400** | 1.82±0.07 | 1.55±0.04 | 1.61±0.04 | 1.55±0.06 |  |
| **14,000** | 1.65±0.03 | 1.53±0.02 | 1.48±0.05 | 1.48±0.07 |  |
| **20,000** | 1.61±0.09 | 1.59±0.09 | 1.51±0.02 | 1.63±0.03 |  |
| **36,000** | 1.63±0.06 | 1.46±0.03 | 1.54±0.08 | 1.53±0.06 |  |
| Dietary Vitamin D Level |  |  |  |  | **0.002** |
| Week |  |  |  |  | 0.65 |
| Vit D x Week |  |  |  |  | 0.69 |

^1^Means ± SEM were given and p-values were generated by repeated measures general linear model with the temporal effect as the repeated measurement and the dietary vitamin D_3_ level as the independent variable. Statistical significance established when p < 0.05 (n=8 hens/dietary treatment).

**Supplementary Table 5.** Calcium (Ca) and phosphorus (P) composition of eggshell, ileal digesta, and feces of 78-wk old Hy-Line Brown laying hens fed diets with different levels of vitamin D_3_ over an eleven-week period^1^

| **Treatment (IU Vitamin D_3_/kg of feed)** | **Eggshell Ca and P (%)** | | **Ileal Digesta Ca and P (%)** | | **^2^Fecal Ca and P (%)** | |
| --- | --- | --- | --- | --- | --- | --- |
|  | **Ca** | **P** | **Ca** | **P** | **Ca** | **P** |
| **400** | 38.1±2.2 | 0.038±0.002 | 3.32±0.84 | 0.773±0.129 | 8.71±0.74^ab^ | 2.01±0.22 |
| **800** | 40.4±0.70 | 0.040±0.001 | 3.94±0.63 | 0.764±0.022 | 8.10±0.55^ab^ | 1.97±0.21 |
| **7,400** | 38.8±1.08 | 0.039±0.001 | 2.51±0.37 | 0.390±0.057 | 9.20±0.30^ab^ | 2.24±0.08 |
| **14,000** | 40.5±0.81 | 0.040±0.001 | 3.76±1.39 | 0.750±0.078 | 10.6±0.83^a^ | 2.18±0.13 |
| **20,000** | 38.8±1.47 | 0.039±0.001 | 3.05±1.14 | 0.834±0.173 | 8.70±0.13^ab^ | 2.08±0.09 |
| **36,000** | 40.7±1.13 | 0.041±0.001 | 2.79±0.37 | 0.728±0.059 | 7.36±0.43^b^ | 2.00±0.10 |
| Diet (**P-Value**) | 0.64 | 0.64 | 0.74 | 0.09 | **0.03** | 0.76 |

^1^Ca and P values were determined by general linear model with dietary vitamin D_3_ as the factor with either Ca or P as predictors and Tukey’s post-hoc test for multiple comparisons. Statistical significance established when p < 0.05. For fecal Ca %, fecal P was used as a predictor and vice versa in the statistical model (n=8 hens/dietary treatment).

^2^Values without a common superscript letter were statistically different.
