## Supplemental Figures for "Dietary super-doses of cholecalciferol fed to aged laying hens illustrates limitation of 24,25-dihydroxycholecalciferol conversion"

### Slide 1
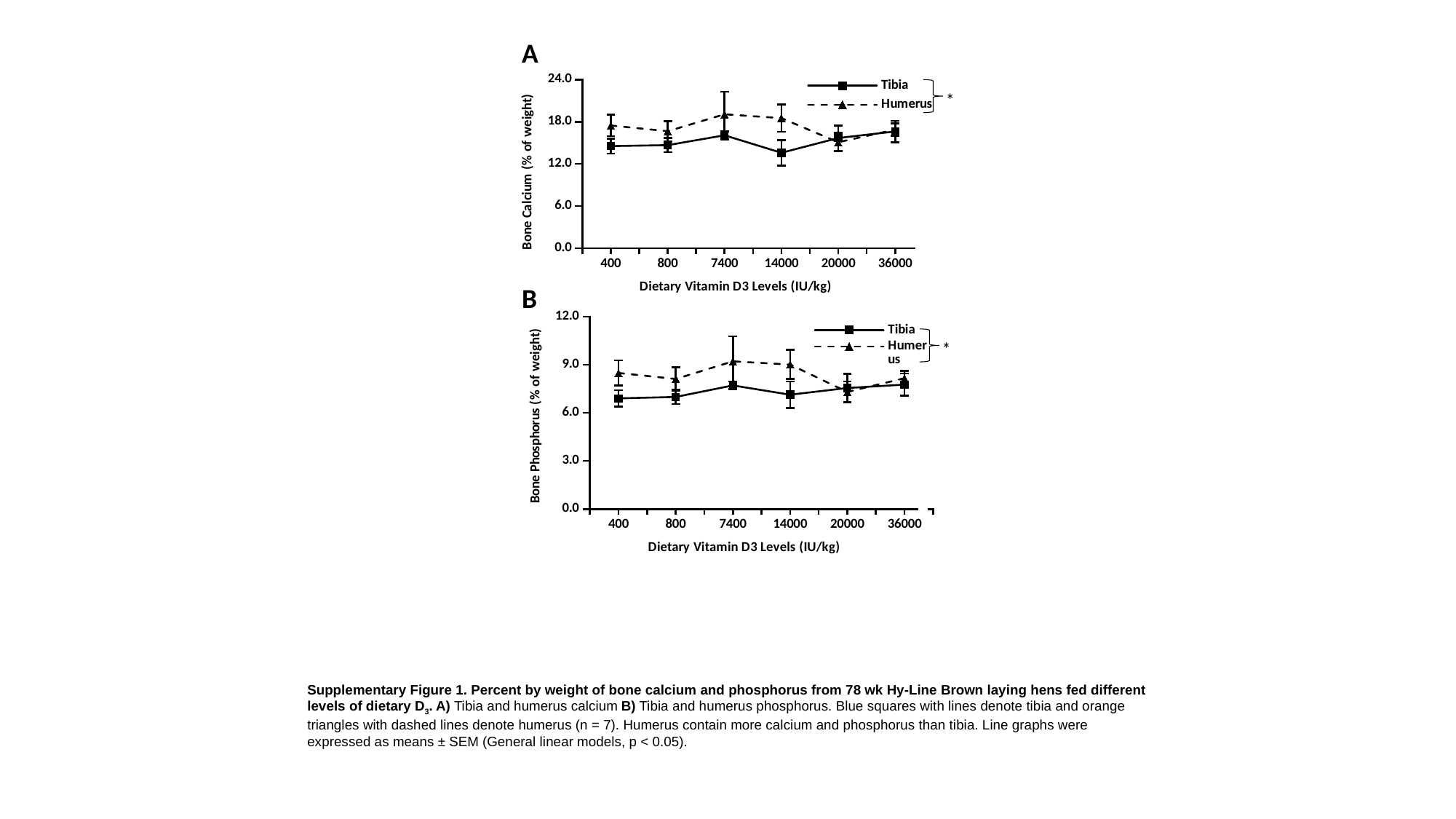

A
#### Chart
| Category | | |
|---|---|---|
| 400 | 14.53291375 | 17.46660375 |
| 800 | 14.670435714285714 | 16.649237 |
| 7400 | 16.072847142857142 | 19.064145 |
| 14000 | 13.577554285714285 | 18.511632857142853 |
| 20000 | 15.682241428571428 | 15.081248333333335 |
| 36000 | 16.61540571428571 | 16.88586857142857 |
*
B
#### Chart
| Category | | |
|---|---|---|
| 400 | 6.904023749999999 | 8.486292500000001 |
| 800 | 6.990613857142857 | 8.112944285714287 |
| 7400 | 7.709115285714285 | 9.212385 |
| 14000 | 7.132385714285713 | 9.010788571428572 |
| 20000 | 7.549435714285714 | 7.301946666666669 |
| 36000 | 7.761444285714286 | 8.164317142857143 |
*
Supplementary Figure 1. Percent by weight of bone calcium and phosphorus from 78 wk Hy-Line Brown laying hens fed different levels of dietary D_3_. A) Tibia and humerus calcium B) Tibia and humerus phosphorus. Blue squares with lines denote tibia and orange triangles with dashed lines denote humerus (n = 7). Humerus contain more calcium and phosphorus than tibia. Line graphs were expressed as means ± SEM (General linear models, p < 0.05).

### Slide 2
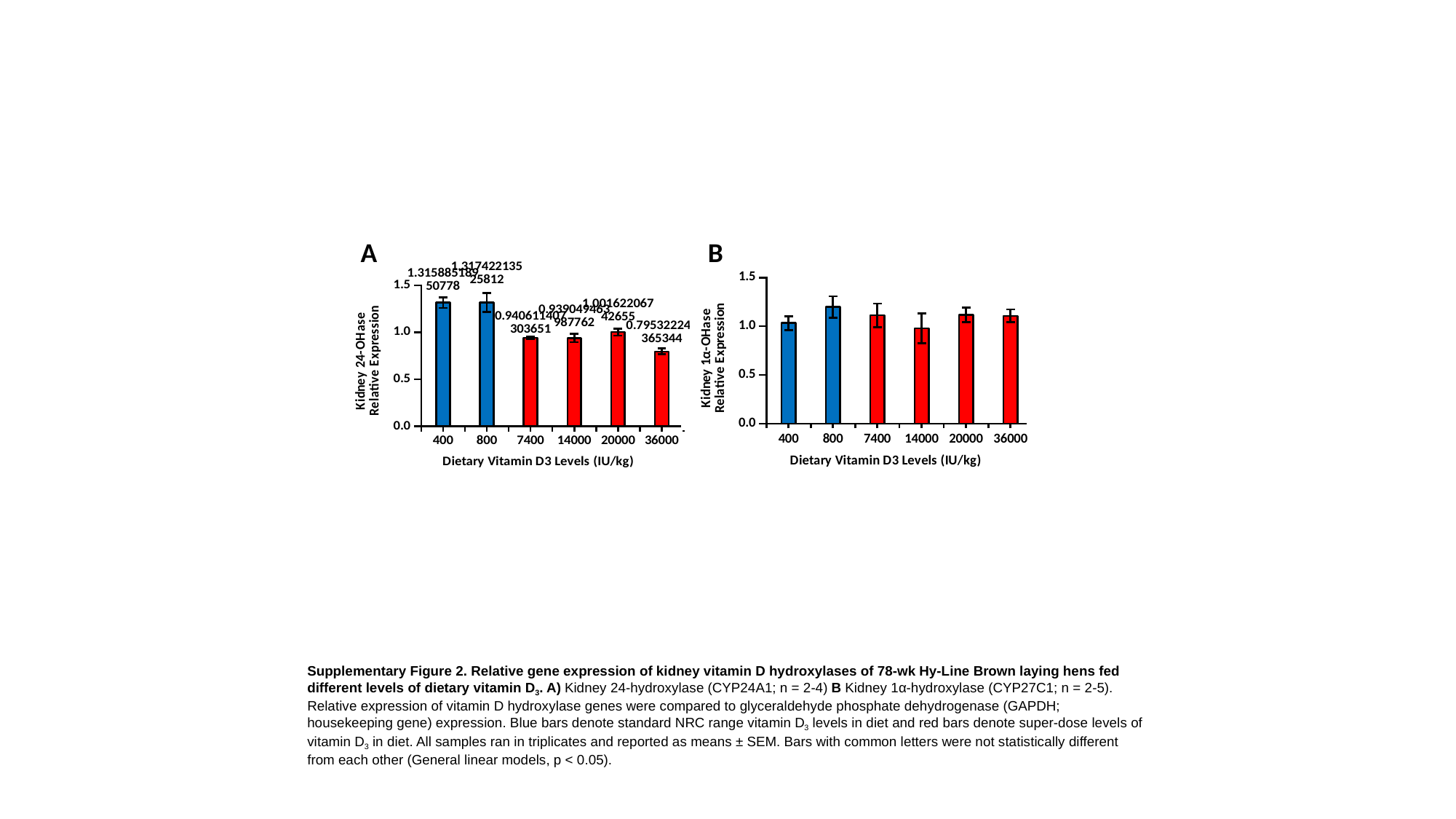

A
B
#### Chart
| Category | Relative Expression |
|---|---|
| 400 | 1.3158851895077834 |
| 800 | 1.3174221352581208 |
| 7400 | 0.9406114073036507 |
| 14000 | 0.9390494639877618 |
| 20000 | 1.0016220674265546 |
| 36000 | 0.7953222463653443 |
#### Chart
| Category | Relative Expression |
|---|---|
| 400 | 1.032673569547871 |
| 800 | 1.1973377183842255 |
| 7400 | 1.1130948724921126 |
| 14000 | 0.9786238540558264 |
| 20000 | 1.117437251545265 |
| 36000 | 1.1068309758319848 |
Supplementary Figure 2. Relative gene expression of kidney vitamin D hydroxylases of 78-wk Hy-Line Brown laying hens fed different levels of dietary vitamin D_3_. A) Kidney 24-hydroxylase (CYP24A1; n = 2-4) B Kidney 1α-hydroxylase (CYP27C1; n = 2-5). Relative expression of vitamin D hydroxylase genes were compared to glyceraldehyde phosphate dehydrogenase (GAPDH; housekeeping gene) expression. Blue bars denote standard NRC range vitamin D_3_ levels in diet and red bars denote super-dose levels of vitamin D_3_ in diet. All samples ran in triplicates and reported as means ± SEM. Bars with common letters were not statistically different from each other (General linear models, p < 0.05).
